# First insights into the *Aurelia aurita* transcriptome response upon manipulation of its microbiome

**DOI:** 10.1101/2023.03.02.530776

**Authors:** Nancy Weiland-Bräuer, Vasiliki Koutsouveli, Daniela Langfeldt, Ruth A. Schmitz

## Abstract

The associated diverse microbiome contributes to the overall fitness of *Aurelia aurita*, particularly to asexual reproduction. However, how *A. aurita* maintains this specific microbiome or reacts to manipulations is unknown. In this report, the response of *A. aurita* to manipulations of its native microbiome was studied by a transcriptomics approach. Microbiome-manipulated polyps were generated by antibiotic treatment and challenging polyps with a non-native, native, and potentially pathogenic bacterium. Total RNA extraction followed by RNAseq resulted in over 155 million reads used for a *de novo* assembly. The transcriptome analysis showed that the antibiotic-induced change and resulting reduction of the microbiome significantly affected the host transcriptome, e.g., genes involved in processes related to immune response and defense mechanisms were highly upregulated. Similarly, manipulating the microbiome by challenging the polyp with a high load of bacteria (2 × 10^7^ cells/polyp) resulted in induced transcription of apoptosis-, defense-, and immune response genes. A second focus was on host-derived quorum sensing interference as a potential defense strategy. Quorum Quenching (QQ) activities and the respective encoding QQ-ORFs of *A. aurita* were identified by functional screening a cDNA-based expression library generated in *Escherichia coli*. Corresponding sequences were identified in the transcriptome assembly. Moreover, gene expression analysis revealed differential expression of QQ genes depending on the treatment, strongly suggesting QQ as an additional defense strategy. Overall, this study allows first insights into *A. aurita’s* response to manipulating its microbiome, thus paving the way for an in-depth analysis of the basal immune system and additional fundamental defense strategies.

## 1. Introduction

Cnidarians, such as the moon jellyfish *Aurelia aurita*, are distributed worldwide and play essential roles in shaping marine ecosystems (Brekhman et al., 2015). Cnidaria are dated back to about 700 million years and are considered a sister group to the Bilateria (Putnam et al., 2007; Park et al., 2012). Thus, they are among the simplest animals at the tissue level organization possessing two germ layers (ectoderm and endoderm) separated by the mesoglea (Ball et al., 2004). In addition to their morphological simplicity, many Cnidaria, particularly Scyphozoa, have a high level of developmental plasticity, allowing for an enormous tolerance, regeneration potential, and asexual proliferation during their life cycle (Richardson et al., 2009). Cnidaria have evolved and are constantly exposed to diverse microorganisms (Liu et al., 2019). This close association with microorganisms has profound effects on various host functions. Recent studies have demonstrated that specific host-associated microbiota can contribute to various host functions. Examples are host metabolism (Ochsenkühn et al., 2017), development (Rook et al., 2017), organ morphogenesis (Sommer and Bäckhed, 2013), pathogen protection and immunity (Moran and Yun, 2015), behavior (Ezenwa et al., 2012), environmental sensing and adaptation (Bang et al., 2018; Ziegler et al., 2019), developmental transitions (Webster and Reusch, 2017; Woznica et al., 2017), and reproduction (Chilton et al., 2015; Jacob et al., 2015).

Evidently, Cnidarians are constantly exposed to microbes in the environment; consequently, molecular analyses have revealed a variety of molecular pathways to respond to microbial exposure (Dierking and Pita, 2020). In the first step, extracellular surface receptors recognize microbe-associated molecular patterns (MAMPs) during microbial epithelium colonization (Chu and Mazmanian, 2013). MAMPs include lipopolysaccharides (LPSs), peptidoglycan (PGN), flagellin, and microbial nucleic acids (Rosenstiel, 2009). In the first line of defense, antimicrobial peptides (AMPs) regulate establishing and maintaining a specific microbiota (Bosch, 2013; Bosch and Zasloff, 2021). Toll-like receptors (TLRs) at the host cell surface further perceive the MAMP signal, initiating MAMP-triggered immunity (Augustin et al., 2010; Bosch, 2013). Downstream of those conserved signaling cascades are stress-responsive transcription factors, including eukaryotic transcription factors of the proteins’ NF-kappaB (NF-kB) family (Zheng et al., 2005). Recent studies revealed that eukaryotic hosts also use quorum quenching (QQ) as a strategy to respond to bacterial colonization (Grandclâment et al., 2016). The hosts interfere with the small molecule-dependent bacterial communication through enzymatic degradation of the autoinducer, blocking autoinducer production, or its reception to control population-dependent behaviors like colonization, biofilm formation, and pathogenesis (Kiran et al., 2017; Mukherjee and Bassler, 2019). In the Cnidarian *Hydra*, the autoinducer signaling molecule 3-oxo-homoserine lactone has been shown to be converted into the inactive 3-hydroxy counterpart by a host-derived oxidoreductase allowing host colonization of the main colonizer *Curvibacter sp*. (Pietschke et al., 2017).

The Cnidarian *A. aurita* harbors a highly diverse and dynamic microbiota specific to the animal, the different sub-populations, and life stages (Weiland-Bräuer et al., 2015a). In the absence of the specific microbial community, the fitness of *A. aurita* was significantly compromised, and notably, asexual reproduction was almost halted (Weiland-Bräuer et al., 2020a). This microbial impact is crucial at the polyp life stage before entering the process of asexual offspring production (strobilation) to ensure a normal progeny output (Jensen et al., 2023). Moreover, in *A. aurita*, three proteins interfering with bacterial QS were identified (Weiland-Bräuer et al., 2019). Incubation of native animals with potentially pathogenic bacteria induced the expression of the identified QQ-ORFs, strongly suggesting a host defense strategy.

Despite the growing knowledge about the impact of microbes on the host and the fundamental strategies of the host to respond, research to understand how microbiomes influence host gene expression is still in its infancy (Nichols and Davenport, 2021). Many studies of model organisms and humans demonstrated an interlinkage between the microbiome and the host’s gene expression. However, the direction of causality mainly remained unanswered (Nichols and Davenport, 2021). Comparing conventional (microbiome-containing) to germ-free systems is one way to assess whether the microbiome plays a causative role in regulating gene expression (Bäckhed et al., 2012; Al-Asmakh and Zadjali, 2015; Fu et al., 2017; Pierre, 2022). Genome-wide transcriptomic analyses are now routinely used to quantify the changing levels of each transcript under different conditions (Conesa et al., 2016).

In the present study, we aimed to determine the influence of the associated microbiota on *A. aurita’s* gene expression. After microbiome manipulation (by antibiotic treatment or bacterial challenge), RNA was extracted from polyps, followed by RNA-Seq and a *de novo* transcriptome assembly. Gene ontology categories and gene expression patterns were analyzed to elucidate how *A. aurita* recognizes and responds to the manipulation of its native microbiome and the presence of potential pathogens. A particular focus was on host-derived QQ activities as an additional potential defense strategy.

## 2. Materials and Methods

### 2.1. *Aurelia aurita* polyp husbandry

Husbandry is described in detail by Weiland-Bräuer *et al*. (Weiland-Bräuer et al., 2015a; Weiland-Bräuer et al., 2020a). Briefly, polyps of the sub-population North Atlantic (Roscoff, France) were kept in the lab in 2-liter plastic tanks in 3 % artificial seawater (ASW) (tropical sea salts; Tropic Marin). Polyps were fed twice a week with freshly hatched *Artemia salina* (HOBBY, Grafschaft-Gelsdorf, Germany).

### 2.2. Reduction of the native *Aurelia aurita* polyp microbiota by antibiotics

Single native polyps were placed in 48 well multiwell plates in 1 mL ASW supplemented with an antibiotic mixture (Provasoli’s antibiotic mixture with final concentrations of 360,000 U/liter penicillin G, 1.5 mg/liter chloramphenicol, 1.8 mg/liter neomycin, and 9,000 U/liter polymyxin B; all components from Carl Roth, Karlsruhe, Germany). No food was provided during the antibiotic treatment. The reduction and consequent change of the microbiota were tested by plating a single homogenized polyp (10 replicates) on Marine Bouillon agar plates (Carl Roth, Karlsruhe, Germany). Plates were incubated for 5 days at 20 °C. Colony forming units (cfu) were calculated, and an 87 ± 9 % reduction per polyp was determined.

### 2.3. Bacterial growth conditions and microbial challenge of polyps

Bacteria (*Pseudoalteromonas espejiana* GenBank accession No. MK967174, and *Vibrio anguillarum* GenBank accession No. MK967055) for the microbial challenge were isolated from *A. aurita* polyps (Weiland-Bräuer et al., 2020b). Strains were grown in Marine Bouillon (MB; Carl Roth, Karlsruhe, Germany) at 30 °C and 120 rpm to optical turbidity at 600 nm of 0.8. *Klebsiella oxytoca* M5aI (DSM No. 7342) was similarly grown in Luria-Bertani (LB) medium. Bacterial cell numbers were determined using a Neubauer count chamber (Assistant, Sondheim vor der Röhn, Germany). Pools of 20 native *A. aurita* polyps were separated in 6-well multiwell plates (Greiner, Kremsmünster, Austria) in 4 mL 3 % ASW after washing them twice with sterile ASW. A pool of 20 native *A. aurita* polyps was incubated with 10^8^ cells/mL (in 4 mL) of the respective strain at 20 °C for 30 min. Next, polyps were washed twice with sterile ASW to remove the bacteria and used to isolate total RNA.

### 2.4. Experimental design for transcriptome analysis

The North Atlantic sub-population polyps were generated from a single mother polyp by clonal budding. Pools of 20 daughter polyps were separated into 6-well plates in 4 ml 3 % ASW. Polyps were kept in five conditions without food supply: 1. native polyps without treatment, 2. antibiotic (AB)-treated polyps, 3. native polyps challenged with 10^8^ cells/mL *Klebsiella oxytoca* M5aI, 4. native polyps challenged with 10^8^ cells/mL *Pseudoalteromonas espejiana*, and 5. native polyps challenged with 10^8^ cells/mL *Vibrio anguillarum* (**Fig. 1**). Three replicates (3 × 20 polyps) were used for each condition. For the bacterial challenges, 20 polyps in 4 mL ASW were supplemented with 4×10^8^ cells and incubated for 30 min (**Fig. 1)**. The total RNA was extracted from each pool of polyps.

**Figure 1:**
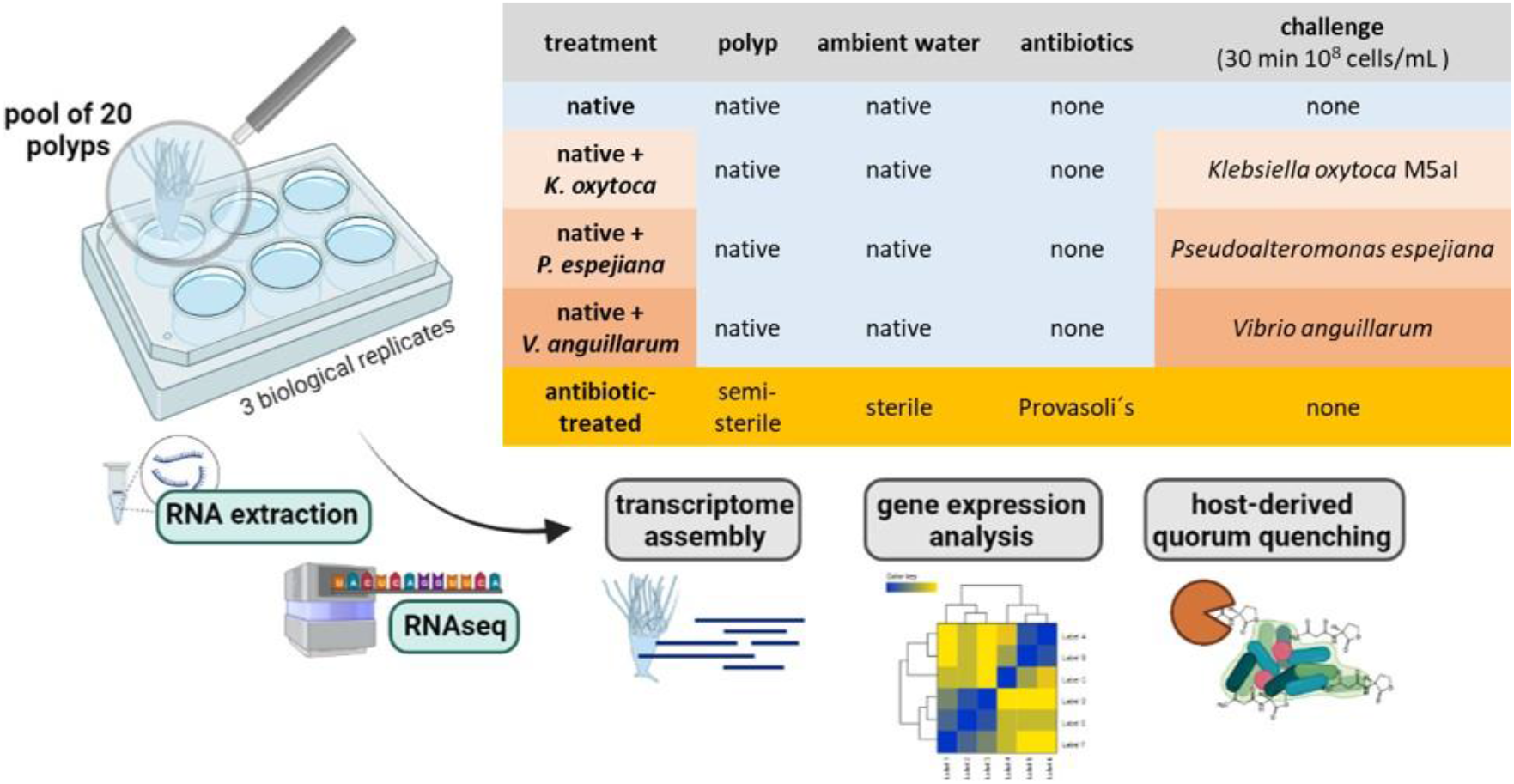
Experimental setup. RNA extraction with subsequent RNAseq and analysis was conducted on a pool of 20 polyps (3 biological replicates). Polyps were kept under native and microbiome-manipulated conditions.

### 2.5. RNA isolation

Total RNA of a pool of 20 *A. aurita* polyps was isolated with an adapted protocol of Gold *et al*., 2019 (Gold et al., 2019). In more detail, polyps were washed three times with sterile ASW (to remove antibiotic residues) and homogenized with a motorized pestle. RiboLock RNase Inhibitor (40 U/μL, Thermo Fisher Scientific, Waltham/Massachusetts, USA), 200 μL lysis solution (100 mM Tris/HCl, pH 5.5, 10 mM disodium EDTA, 0.1 M NaCl, 1 % SDS, 1 % ß-mercaptoethanol), and 2 μl Proteinase K (25 mg/mL, Thermo Fisher Scientific, Waltham/Massachusetts, USA) were added to the homogenate and incubated for 10 min at 55 °C. Chilled solutions were added with 5 μL of 3 M sodium acetate (pH 5.2) and 250 μL phenol-chloroform-isoamyl alcohol (25:24:1) and incubated for 15 min on ice prior to 15 min centrifugation at 12,000 x g at 4 °C. The upper phase was mixed with 1 volume 2-propanol, and precipitation occurred at -80 °C overnight. The precipitate was centrifuged for 15 min at 12,000 x g at 4 °C. The pellet was washed twice with 70 % ethanol before the air-dried pellet was dissolved in 25 μL RNase-free water. DNA contaminations were removed with Turbo DNA-free DNase (Thermo Fisher Scientific, Waltham/Massachusetts, USA). The RNA quality and quantity were assessed by NanoDrop1000 (Thermo Fisher Scientific, Waltham/Massachusetts, USA) and 1.5 % agarose gel electrophoresis, and the cDNA library was prepared with DNA-free host RNA (500 ng) using the TruSeq Stranded mRNA Library Preparation kit. The library was sequenced with the NextSeq 500 System (Illumina, San Diego/ California, USA).

## 2.6. Transcriptome analysis

The raw reads were trimmed with Trimmomatic (Bolger et al., 2014) to remove bad-quality reads and the adapters. A *de novo* assembly with the trimmed sequences was conducted with the Trinity package v2.8.4 (Grabherr et al., 2011b; a). The quality of the *de novo* assembly was assessed with several parameters, such as the N50 value and the percentage of the assembly-mapped reads, which was calculated with Bowtie2. Finally, Benchmarking Universal Single-Copy Orthologs (BUSCO V2/3) against metazoan cassettes (Simão et al., 2015) was used to evaluate the completeness of the assembly regarding the core genes found in metazoans.

A Blastx of the transcriptome assembly was executed against the Swiss-Prot database for metazoans. For the gene expression analysis, reads were mapped to the reference assembly with Bowtie2 (Langmead and Salzberg, 2012). Transcript quantification was performed with RSEM (Li and Dewey, 2011). The differential gene expression (DGE) analysis was done with edgeR (Robinson et al., 2010; McCarthy et al., 2012). Pairwise comparisons between all experimental conditions were performed using FDR ≤ 0.001 and 2-fold changes as statistical parameters. Furthermore, GO enrichment analysis was conducted using a Fisher’s Exact Test in Blast2GOPRO (Conesa et al., 2005) with a p-value threshold of ≤ 0.05. Here, the total annotation file of the reference transcriptome was used as the “reference dataset“, whereas the upregulated genes in each condition served as the “test dataset“. Based on sequence depth and quality, only two out of three biological replicates were used for the gene expression analysis for the conditions of native polyps and native polyps under the bacterial challenges. Transcriptome data is deposited under the BioProject ID PRJNA938117.

### 2.7. Quorum quenching (QQ) assay

QQ assays using the reporter strains AI1-QQ.1 and AI2-QQ.1 were performed with cell-free supernatants and cell extracts of EST clones from the *A. aurita* EST library as described (Weiland-Bräuer et al., 2015b). Following the manufacturer’s protocol, the plasmids of identified QQ-active single EST clones were purified using the Presto Mini Plasmid kit (GeneAid, New Taipeh City, Taiwan). The respective inserts were Sanger sequenced at the Institute of Clinical Molecular Biology in Kiel with the primer set T7_Promoter (5′-TAATACGACTCACTATAGGG-3′) and T7_Reverse (5’-TAGTTATTGCTCAGCGGTGG-3’); submission ID 2678799.

## 3. Results

A transcriptomics approach was applied to gain insights into the response of *A. aurita* polyps to manipulation of its native, associated microbiota, with a particular focus on quorum sensing interference as a potential host defense strategy. In general, pools of 20 polyps were treated and processed together in the experiments, each with three biological replicates using the following treatments: native, antibiotic-treated polyps resulting in 87 ± 9 % reduced microbial cells per polyp (cfu/polyp), and native polyps challenged with *Klebsiella oxytoca* M5aI, *Pseudoalteromonas espejiana*, or *Vibrio anguillarum* (2 × 10^7^ cells/polyp, **Fig. 1**). The respective pools were used for total RNA extraction followed by RNAseq and transcriptome analysis.

### 3.1. Statistics of the *de novo* transcriptome assembly

The RNAseq approach overall resulted in 167,269,257 raw reads. After quality trimming, 145,746,530 reads (87.1 % of the total reads) remained for further analysis and were used for the *de novo* transcriptome assembly (**Tab. S1A**). The *de novo* assembly resulted in 213,897 transcripts and 160,700 genes (**Tab. S1A**) with an N50 value of 1,170 (**Tab. S1B**). 94.25 % of reads were successfully aligned back to the assembly, while the completeness of genes was 96.42 % according to the BUSCO score for metazoans (**Tab. S1B**). 30 % (597,796 transcripts) of the assembly revealed an annotation against the Swiss-Prot database for metazoans (**Tab. S2**).

### 3.2. Manipulation of its microbiome affects *A. aurita’s* transcriptome

Transcriptome analysis identified five separated clusters corresponding to the different conditions, while biological replicates of a condition cluster together (**Fig. 2**). It should be noted that three replicates were only analyzed for antibiotic (AB)-treated polyps, as all other conditions resulted in an unacceptable low sequence depth for one replicate. Notably, two clusters were identified within the hierarchical clustering of transcripts. Here, native and AB-treated conditions yielded a tremendous difference. AB-treated polyps showed a massive reduction of the microbial load (by 87 ± 9 %), assuming a rigorous change in the abundance and diversity of microbial colonizers. Bacteria-challenged polyps showed a mixture of native and AB-treated expression patterns (**Fig. 2**). In more detail, the comparison of native and AB-treated polyps resulted in the highest number of differentially expressed genes (22,073 genes), with 10,451 upregulated and 11,622 down-regulated genes in AB-treated compared to native polyps (**Tab. S3A**). According to the GO enrichment analysis (Conesa et al., 2005), the 10,451 upregulated genes were overrepresented in processes related to inflammation and immune response (e.g., acute inflammatory response to antigenic stimuli, MAP-kinase activity, cytokine production, and neutrophil cell activation) (**Fig. 3A; Tab. S4A**). Genes involved in immune response and apoptosis included, e.g., Caspases, Apoptosis regulators, Interferones, and Toll-like receptors (**Fig. 3B, Tabs. S3A, S4A**). On the contrary, down-regulated genes in AB-treated polyps were enriched for processes related to development, morphogenesis, reproduction, stimuli response, and signaling (**Figs. 3C, D; Tabs. S3A, S4A**).

**Figure 2:**
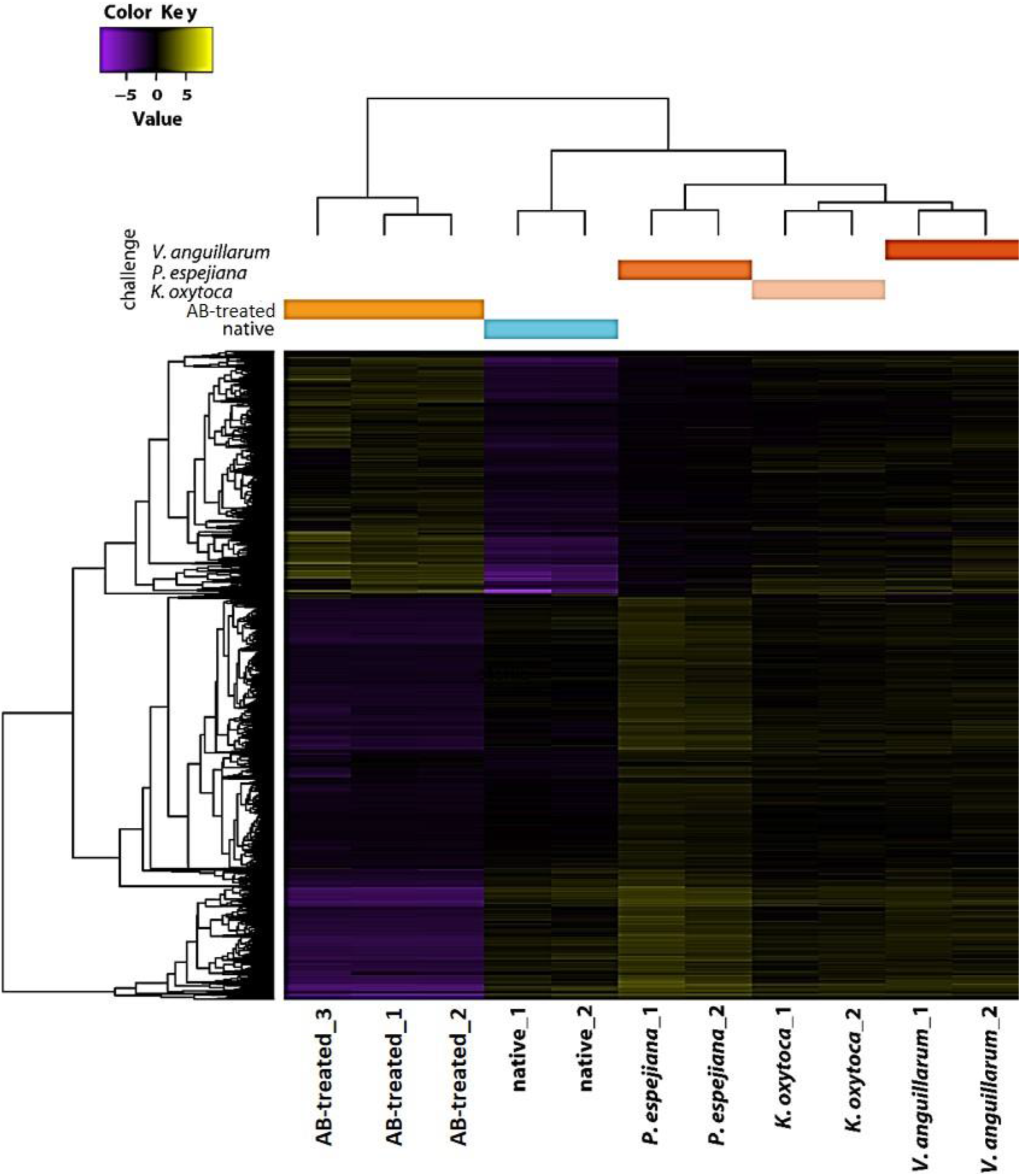
Holistic expression patterns of *A. aurita* polyps. Heatmap including the Differential Expressed (DE) genes in all the pairwise comparisons of the native, Ab-treated, and bacteria-challenged polyps, including replicates.

**Figure 3:**
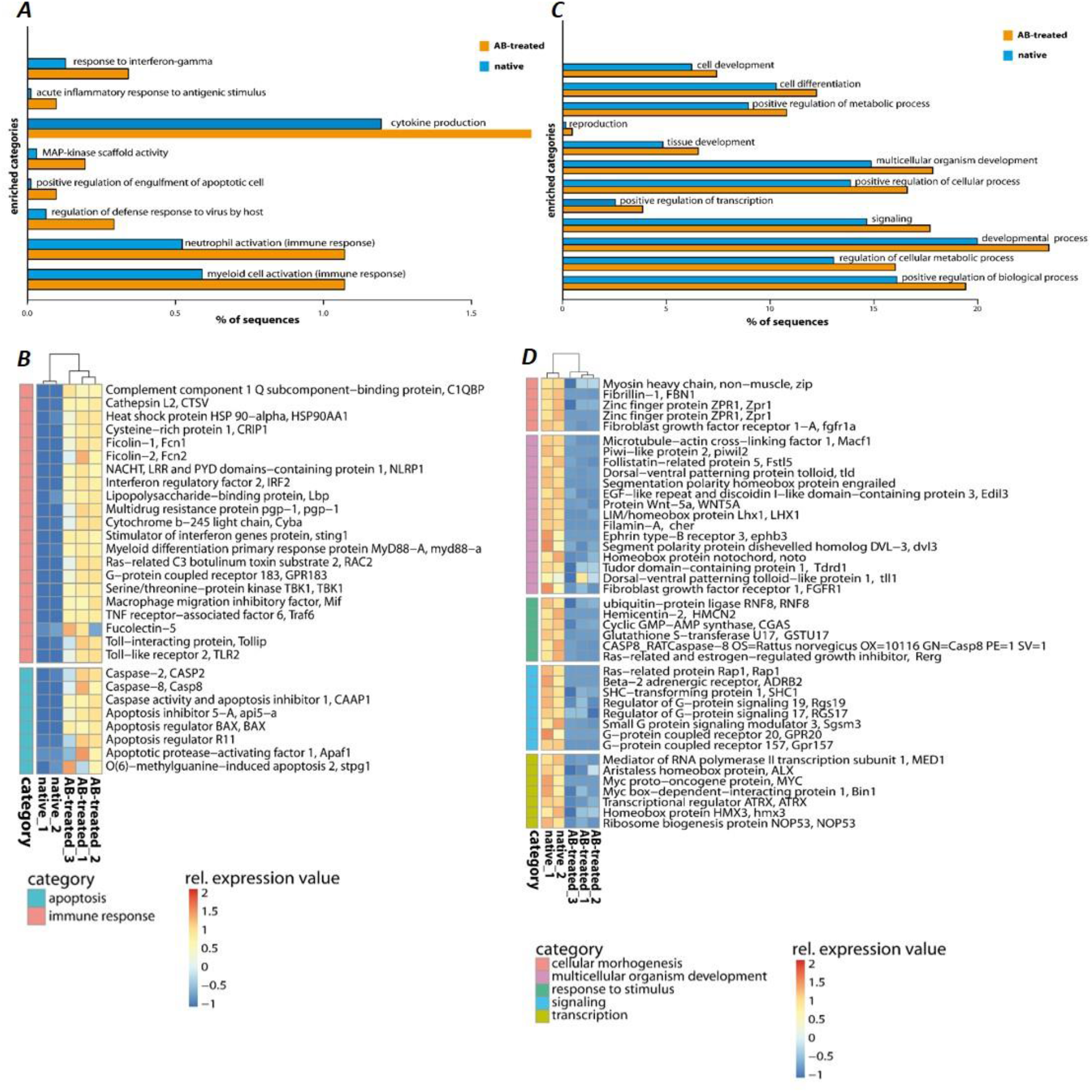
GO enrichment analysis of differentially expressed genes of native and Ab-treated *A. aurita* polyps. ***(A, C)*** Barplots with GO-enriched gene categories when comparing native and Ab-treated polyps. Barplots indicate the proportion (%) of DE gene sequences involved in ***(A)*** upregulated and ***(C)*** down-regulated GO-enriched categories compared to the reference (*de novo* assembly). ***(B, D)*** Heatmap of upregulated and down-regulated genes with the comparison of native and AB-treated polyps related to ***(B)*** immune response and apoptosis ***(D)*** and common physiological processes. Relative expression level increase from blue to red.

When comparing expression patterns between native and bacteria-challenged polyps, differences based on the bacterial species used in each challenge were revealed (**Fig. 4**). The comparison of native polyps compared to those challenged with *V. anguillarum* gave the highest number of differentially expressed (DE) genes (7,193 genes; **Fig. 4A, Tab. S3B**), followed by polyps challenged with *K. oxytoca* (6,974 DE genes; **Fig. 4A, Tab. S3C**), and *P. espejiana* (2,721 genes; **Figs. 4A, Tab. S3D**). We observed 2,210 genes were commonly upregulated among the treatments compared to native polyps (**Fig. 4A, Tab. S3E**). At the same time, 52 genes were jointly downregulated (**Fig. 4B, Tab. S3E**). Genes related to biological processes, such as defense, immune and inflammatory responses, and the regulation of apoptotic processes, were upregulated in bacteria-challenged polyps (**Fig. 4B, Tab. S3E**). In contrast, genes related to organism development, cell cycle processes, and cytoskeleton organization were down-regulated in challenged polyps (**Fig. 4B, Tab. S3E**). Moreover, exclusively upregulated and down-regulated genes were identified for each bacterial challenge (**Fig. 4)**. 1,319 genes were exclusively upregulated in *V. anguillarum*-challenged polyps (**Fig. 4A left panel, Tab. S3F**), 1,020 genes were exclusively upregulated in polyps challenged with *K. oxytoca* (**Fig. 4A left panel, Tab. S3G**), and 160 genes exclusively upregulated in *P. espejiana*-challenged polyps (**Fig. 4A left panel, Tab. S3H**).

**Figure 4:**
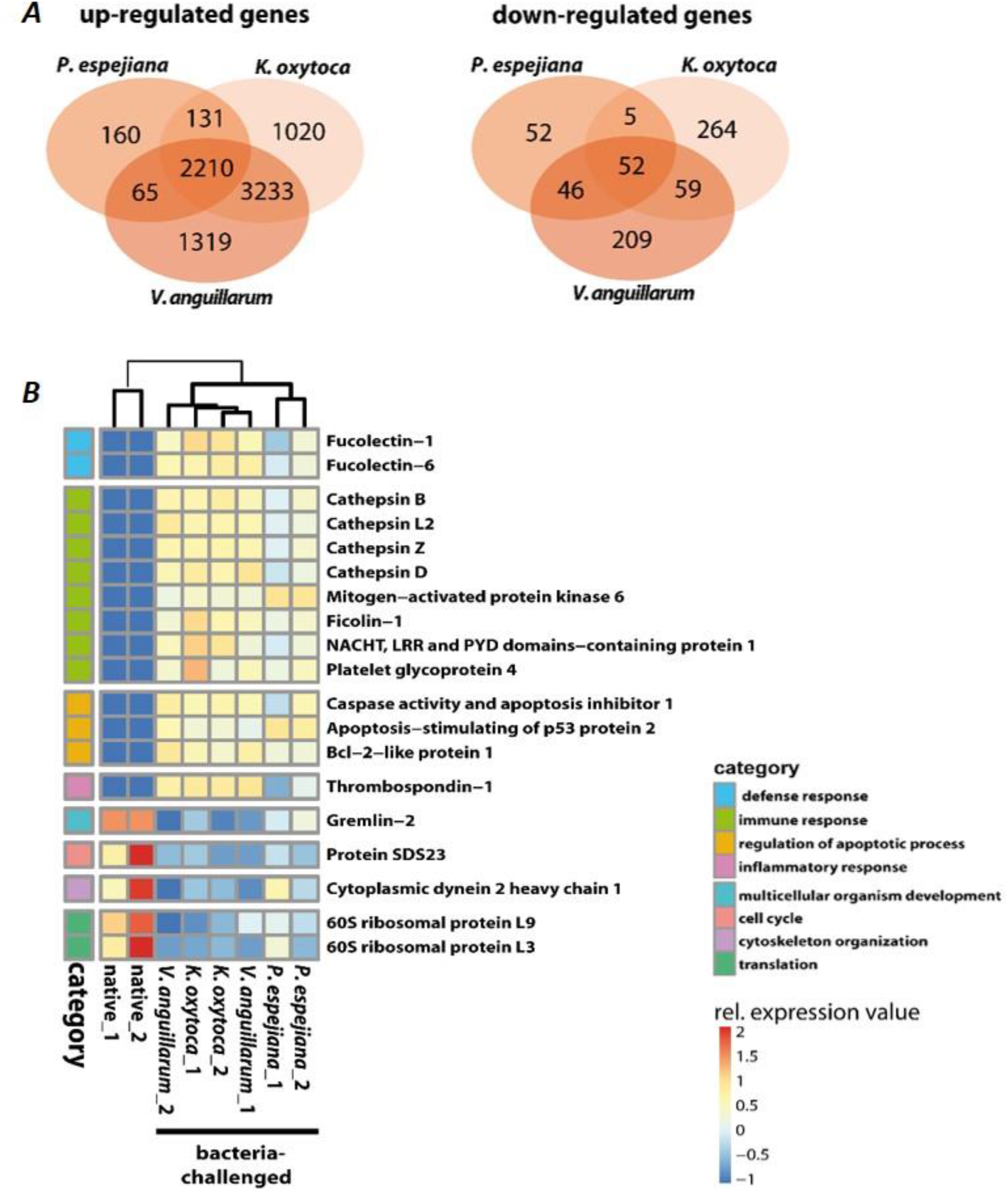
GO enrichment and differential expression analysis of genes divergently transcribed in native compared to bacteria-challenged polyps. ***(A)*** Venn diagram indicating the genes that are commonly and exclusively upregulated (left panel) or down-regulated (right panel) in the different bacteria-challenged conditions. ***(B)*** Heatmap of up- and downregulated genes in native compared to bacteria-challenged polyps. Relative expression level increases from blue to red.

Similarly, exclusively down-regulated genes were identified (**Fig. 4A, right panel**). Analyzing the GO-enriched categories for each species (**Tab. S4B-D**), we found categories common among all treatments (**Tab. S4E**) as well as categories present in two different treatments or only in one (**Fig. 4B, Tab. S4B-D**). Focusing on genes exclusively upregulated after each species-specific challenge, we indeed observed that the same GO categories, e.g., immune response, MAPK signal transduction, autophagy, and DNA repair, were enriched to different degrees, though represented by various genes (depicted in **Fig. 5**).

**Figure 5:**
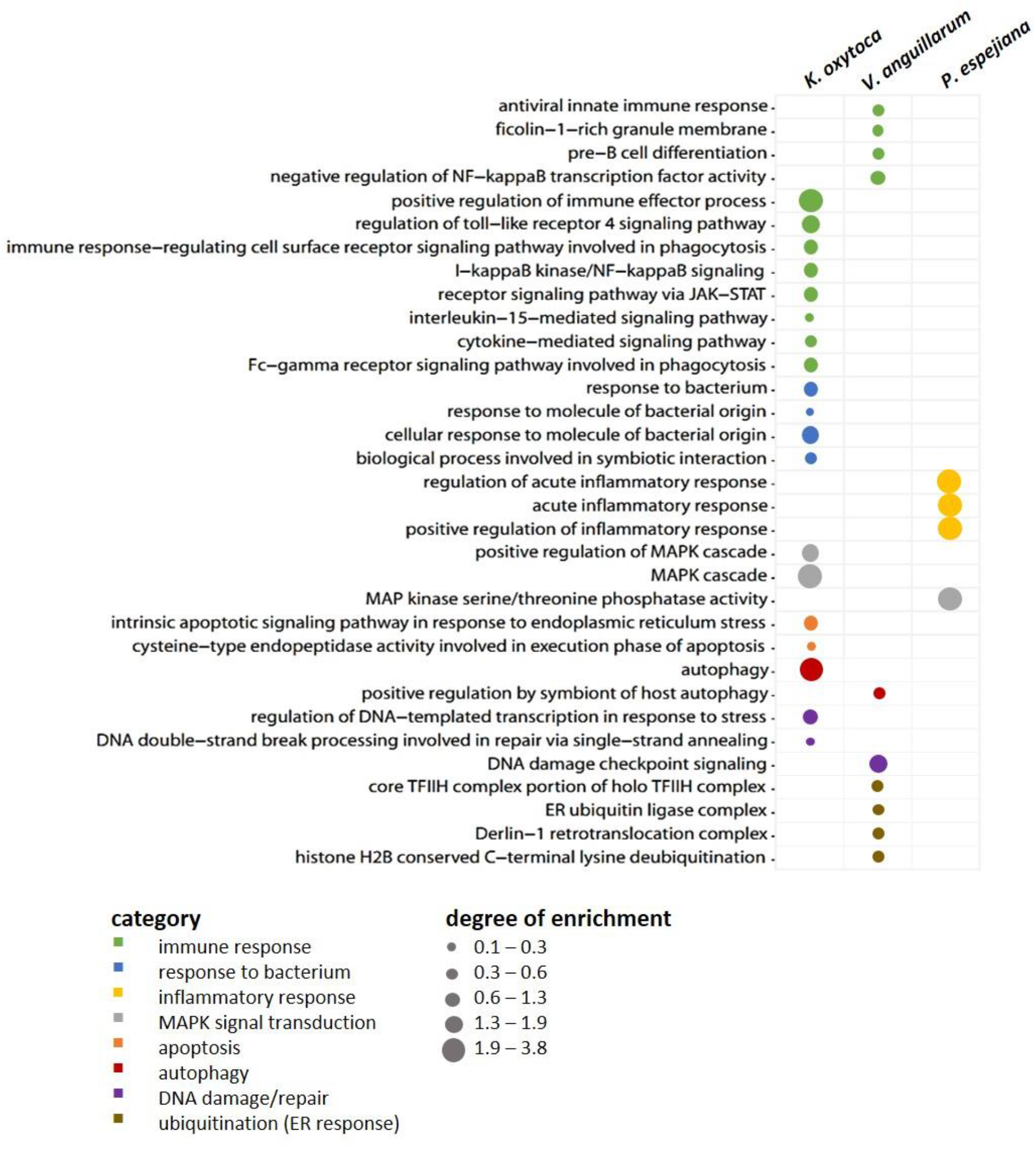
Exclusively upregulated genes in native vs. bacteria-challenged polyps. Bubble plot showing the degree of enrichment per category of exclusively upregulated genes in each bacteria-challenged compared to native conditions.

### 3.3. Bacterial challenge of polyps affects host Quorum quenching

Recent studies revealed that interfering with Qurom sensing, so-called quorum quenching (QQ), might be a fundamental, additional inter-phylum interaction to maintain metaorganismal homeostasis. Consequently, we aimed to identify host-derived QQ activities. An expressed sequence tag (EST) library from *A. aurita* polyps-derived mRNA was constructed in *E. coli* SoluBL21 (Ladewig et al., 2023). The library consisted of 29,952 clones with an insertion efficiency of approx. 98 % and an average insert size of 1.46 kbp, resulting in 43 Mbps cloned host transcriptome, corresponding to an estimated 11.4 % coverage (calculated *A. aurita* genome size 376 Mbps (Gold et al., 2019)). The *A. aurita* EST library was successively screened for QQ activities towards acyl-homoserine lactones (AHL) and autoinducer-2 (AI-2) in cell-free cell extracts and culture supernatants of the EST clones using established *E. coli*-based reporter systems (Weiland-Bräuer et al., 2015b). Overall, 37 out of 29,952 EST clones were identified as QQ active (**Tab. 1**). Predominantly, QQ activities against the Gram-negative signaling molecule AHL were detected. Although those host-derived QQ activities were identified in a functional screen, their biological activities *in vivo* and their functions in *A. aurita* have to be explored. In a first attempt, plasmid insertions of QQ-active single EST clones were sequenced, and sequence data subsequently checked for homologies in the *A. aurita* transcriptome assembly. All QQ-conferring sequences were identified in the transcriptome assembly with homologies ranging from 83 to 100 % (**Tab. S5**). Furthermore, public databases (NCBI, UniProt, PFAM) were used for homology predictions and annotation (**Tab. S5**). Unexpectedly, several QQ-ORFs showed homologies to highly conserved ribosomal proteins, suggesting a moonlighting function of those proteins (Jeffery, 2003; Singh and Bhalla, 2020). Secondly, analyzing transcription levels of those identified QQ-ORFs in the generated RNAseq data set (see above) allowed first insights into their transcriptional regulation in response to the respective treatments, which might support predicting their biological role. Relative expression of the 37 QQ-conferring genes was calculated for microbiome-manipulated versus native polyps. Hierarchical clustering observed four clusters of expression profiles. The first cluster represents QQ genes similarly expressed in microbiome-manipulated polyps compared to native ones (**Fig. 6**, right column highlighted in grey). The second cluster of genes showed decreased expression in all treatments, except when challenged with *P. espejiana* (**Fig. 6**, right column light orange cluster). The third cluster summarizes those genes with increased expression regardless of the type of manipulation (**Fig. 6**, right column dark orange cluster). The fourth cluster includes those genes with an increased expression after adding potential pathogens but reduced expression when polyps were AB-treated (**Fig. 6**, right column green cluster). Notably, *P. espejiana* primarily showed increased expression of QQ-ORFs; thus, hierarchal clustering showed the highest dissimilarity compared to the other treatments.

**Table 1:** *Aurelia aurita*-derived quorum quenching activities derived from an expressed sequence tag (EST) library. Functionally identified QQ-ORFs are listed with their QQ activity in the cell-free cell extract (CE) and culture supernatant (SN) against acyl-homoserine lactones (AHL) and autoinducer-2 (AI-2) with their potential annotations; x, activity; -, no activity.

| QQ-ORF | Original clone designation | QQ activity |  |
| --- | --- | --- | --- |
|  |  | AHL | AI-2 |
| QQ_Aa_1 | 115/H7 | x | x |
| QQ_Aa_2 | 118/E4 | x | - |
| QQ_Aa_3 | 118/G4 | x | x |
| QQ_Aa_4 | 127/C7 | x | x |
| QQ_Aa_5 | 164/A6 | x | - |
| QQ_Aa_6 | 164/E2 | x | x |
| QQ_Aa_7 | 184/A11 | x | - |
| QQ_Aa_8 | 184/B9 | x | - |
| QQ_Aa_9 | 202/G6 | x | - |
| QQ_Aa_10 | 208/E9 | x | - |
| QQ_Aa_11 | 208/A3 | x | - |
| QQ_Aa_12 | 213/E2 | x | - |
| QQ_Aa_13 | 213/F2 | x | - |
| QQ_Aa_14 | 217/H4 | x | - |
| QQ_Aa_15 | 221/C11 | x | x |
| QQ_Aa_16 | 284/H11 | x | - |
| QQ_Aa_17 | 115/A1 | x | - |
| QQ_Aa_18 | 115/A11 | x | x |
| QQ_Aa_19 | 115/G1 | x | - |
|  |  | AHL | AI-2 |
| QQ_Aa_20 | 118/F5 | x | x |
| QQ_Aa_21 | 164/B2 | x | - |
| QQ_Aa_22 | 164/C8 | x | - |
| QQ_Aa_23 | 164/D1 | x | - |
| QQ_Aa_24 | 164/D6 | x | - |
| QQ_Aa_25 | 164/F2 | x | - |
| QQ_Aa_26 | 164/G1 | x | - |
| QQ_Aa_27 | 127/A8 | x | x |
| QQ_Aa_28 | 127/A10 | x | - |
| QQ_Aa_30 | 127/C12 | x | - |
| QQ_Aa_31 | 127/F8 | x | - |
| QQ_Aa_32 | 184/A6 | x | - |
| QQ_Aa_33 | 213/B7 | x | - |
| QQ_Aa_34 | 223/E7 | x | - |
| QQ_Aa_35 | 270/C5 | x | x |
| QQ_Aa_36 | 273/A1 | x | - |
| QQ_Aa_37 | 273/F12 | x | x |
| QQ_Aa_38 | 284/A10 | x | x |

**Figure 6:**
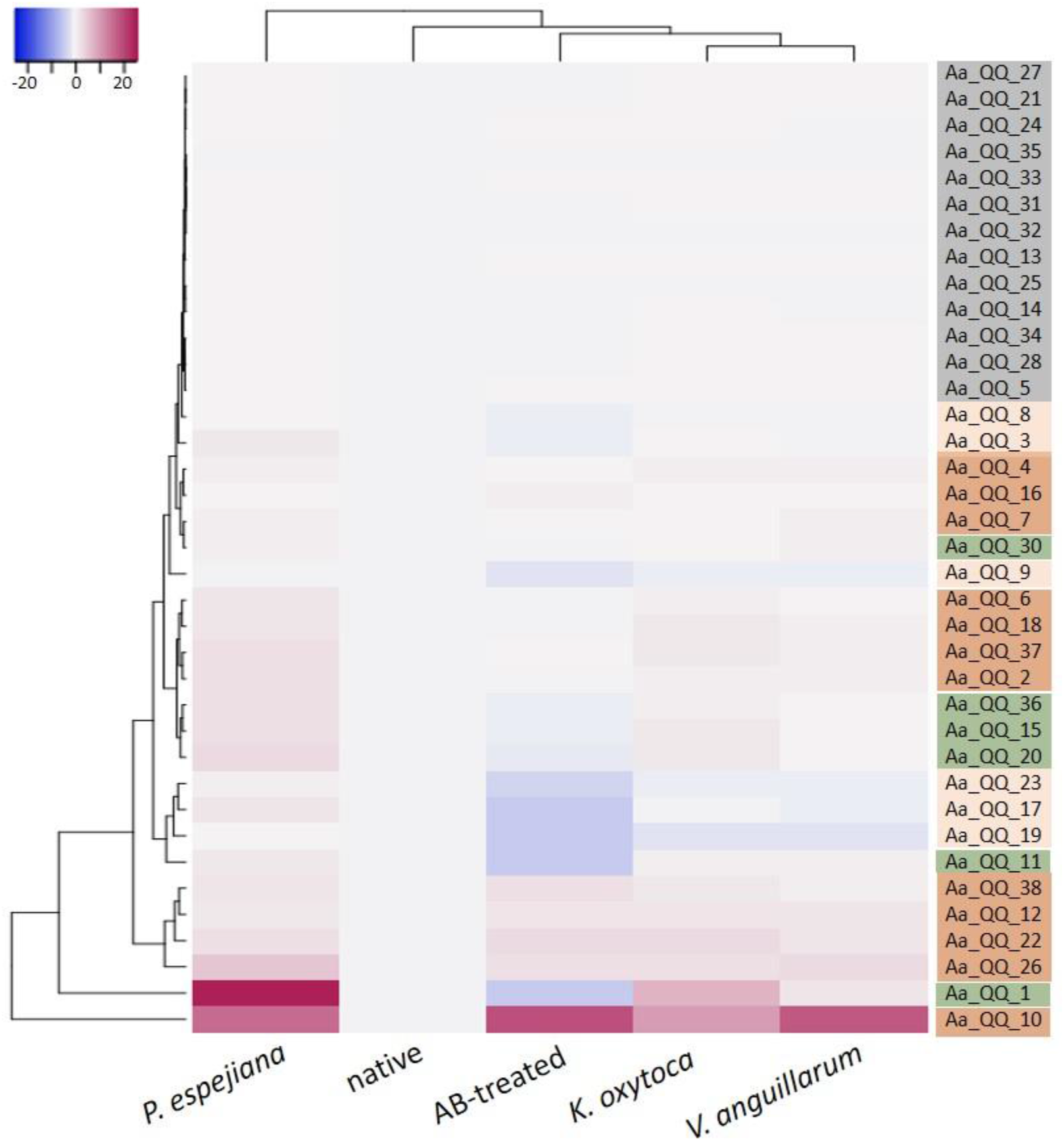
Differential expression of functionally identified *A. aurita* QQ-ORFs. Heatmap visualizes the expression patterns of 37 identified host-derived QQ-ORFs per condition as a mean of 2 to 3 replicates in combination with hierarchical clustering of conditions and expression profiles of ORFs.

## 4. Discussion

In the present study, we obtained the first insights into the transcriptomic response of *A. aurita* to microbiome manipulation. The manipulations of the polyp microbiome included a massive reduction of viable bacterial cells (87 % reduction) due to applying a broad-spectrum antibiotics mixture. The mixture contained Penicillin, Polymyxin B, Chloramphenicol, and Neomycin

(Provasoli and Pintner, 1980; Liu et al., 2017a; KleinJan et al., 2022), and has been shown to only partially eliminate an entire bacterial community (Azma et al., 2010; Liu et al., 2017a). We assume that by drastically reducing the colony-forming units by 87 %, the diversity and abundance of community members on the polyp are crucially changed. Consequently, the polyp was confronted with the loss of certain community members and a drastic change in the relative abundance of remaining ones. This extreme change in microbiome composition resulted in the up-regulation of various immune response mechanisms. Pattern recognition receptors (PRRs) like Toll-like receptors and Ficolins were upregulated, inducing the first line of defense (**Fig. 3**). Further, Caspases were enriched in AB-treated polyps, potentially maintaining homeostasis through regulating cell death and inflammation (McIlwain et al., 2013). Contrarily, all processes involved in morphogenesis and development, thus not essential for survival, were down-regulated in AB-treated polyps (**Fig. 3**).

Furthermore, the native microbiome was affected by challenging the polyps with three different bacteria in high cell numbers: (i) *K. oxytoca* has not been detected in any life stage of *A. aurita* and thus represents a non-native bacterium (Weiland-Bräuer et al., 2015a; Weiland-Bräuer et al., 2019); (ii) *V. anguillarum*, an opportunistic pathogen of various invertebrates and vertebrates, has been isolated from an *A. aurita* polyp (Austin, 2010; Frans et al., 2011; Weiland-Bräuer et al., 2015a; Weiland-Bräuer et al., 2020b); and (iii) *P. espejiana* was initially isolated from seawater but was also found in high abundance associated with all life stages of *A. aurita*, thus representing a native bacterium (Isnansetyo and Kamei, 2009; Weiland-Bräuer et al., 2015a; Weiland-Bräuer et al., 2020b). Due to bacterial challenges, we observed the up-regulation of defense, immune, and inflammatory responses, and apoptosis regardless of the bacterial species. Based on this finding, we hypothesize that the host recognized the non-native and native bacteria as a potential threat, at least when present in such high cell numbers. The immune system of *A. aurita* likely responded with the four primary innate immune system functions as demonstrated for other Cnidarians and already partly observed for AB-treated polyps (Miller et al., 2007; Parisi et al., 2020). First, immune recognition occurred by PRRs, like Fucolectins, Ficolins, and NACHT-containing domain proteins, binding to bacterial MAMPs/PAMPs (**Figs. 4, 5**). Subsequently, various transcription factors from the NF-κB family were activated (**Figs. 4, 5**). Intracellular signaling cascades (MAPK, interleukin, cytokines) led to target gene transcription to eliminate the threat and mitigate self-harm (**Figs. 4, 5**). Lastly, autophagy, DNA repair, and programmed cell death were upregulated after bacterial challenge (**Figs. 4, 5**). Our observations are consistent with previous results showing that *A. aurita*’s general fitness and, in particular, its asexual reproduction were drastically affected in the absence of microbes and due to the manipulation of the microbiota by challenging with non-native colonizers, e.g., *P. espejiana* and *V. anguillarum* (Weiland-Bräuer et al., 2020a).

The acquisition, establishment, and maintenance of a specific microbiota are advantageous for the host, and its disturbance can contribute to developing diseases (Zheng et al., 2020). The host strives to maintain this homeostasis and presumably uses other defense mechanisms besides the innate immune system (Nichols and Davenport, 2021). The use of bacterial communication and its interference by the host has been recently regarded as an additional interaction mechanism within the complex interplay of the host and its microbiome (White et al., 2020; Weiland-Bräuer, 2021). Several studies evaluated the implication of highly conserved paraoxonases (PON1, PON2, and PON3) as QQ enzymes of the host defense against the pathogen *Pseudomonas aeruginosa* (Mochizuki et al., 1998; Draganov et al., 2005; Stoltz et al., 2007). In the present study, we functionally detected QQ activities of *A. aurita* in a cDNA expression library and verified the respective transcripts within the *de novo* assembled host transcriptome. Their native expression in response to microbiome manipulation resulted in four different expression profiles (**Fig. 6**). A first cluster represents QQ-ORFs similarly low expressed in treated and native polyps. Those findings argue against a biological function in the defense against pathogens. The second cluster showed decreased expression in all treatments except when challenged with *P. espejiana*. Here, a specific defense against *P. espejiana* can be assumed. This assumption aligns with previous studies showing massive declined survival rates in the presence of this bacterium (Weiland-Bräuer et al., 2020a). The multitude of initiated defense mechanisms in the presence of the ubiquitous, native *P. espejiana* (various QQ-ORFs and acute inflammatory response, **Figs. 5, 6**) might indicate that this bacterium represents a threat to *A. aurita*, at least in the high cell numbers, and must be rapidly defended. The third type of expression profile showed increased expression regardless of the type of microbiome manipulation, assuming a general defense. Among those is Ferritin, generally regarded as an intracellular iron storage protein and suggested to be involved in response to infection in different organisms, including fish and marine invertebrates (Moreira et al., 2020). The expression of Ferritin or Ferritin-homologs was upregulated in different tissues in response to bacterial infections or stimulation with LPS (Neves et al., 2009; He et al., 2013; Ren et al., 2014; Chen et al., 2016; Sun et al., 2016; Liu et al., 2017b; Martínez et al., 2017). Furthermore, Ferritin was shown to protect shrimp and fish from viral infections (Ye et al., 2015; Chen et al., 2018). The fourth cluster was expressed in the presence of the non-native and native bacteria, regarded as potential pathogens, but not after antibiotic treatment, assuming a fast response to potential pathogens’ colonization possessing similar PAMPs. Homologies to already described genes within the NCBI database of those QQ-ORFs to predict their function were rare. QQ-ORFs Aa_QQ_15 and Aa_QQ_30, belonging to the fourth cluster, showed non-significant homologies to a chitinase and acetyl-transferase, respectively. Here, enzymatic modification (hydrolysis or acetylation) of the bacterial signaling molecule can be speculated. The four other QQ-ORFs within this expression profile showed the best homologies with highly conserved proteins, including actin, heat shock, and ribosomal proteins. Particularly, ribosomal proteins crucially involved in translation have been shown to comprise other functions (Hurtado-Rios et al., 2022), thus recognized as moonlighting proteins. Moonlighting proteins are capable of performing more than one biochemical function within the same polypeptide chain, including inhibiting infectious bacteria, viruses, parasites, fungi, and tumor cells (Jeffery, 2003; Gao and Hardwidge, 2011; Wang et al., 2015; Singh and Bhalla, 2020). Thus, they have been considered antimicrobial peptides (AMPs) (Hurtado-Rios et al., 2022). Consequently, we hypothesize that the identified QQ activity, besides their canonical function, is promiscuous. Future studies have to be conducted to verify the biological function of the identified *A. aurita* QQ-ORFs, e.g., by genetic and biochemical approaches.

In conclusion, this study provides the first *A. aurita* transcriptome analysis focusing on the impact of microbiome disturbance on host gene expression. Overall it highlights the importance of the native microbiome since processes like morphogenesis, development, and response reactions are down-regulated when it is disturbed due to antibiotics. Microbiome disturbance further induced apoptosis and immune responses, indicating the microbiome’s protective function. Similarly, challenging *A. aurita* with high loads of bacteria resulted in the up-regulation of defense, immune, and inflammatory responses to maintain metaorganismal homeostasis. We have further received indications that quorum quenching is likely an additional mechanism for maintaining the specific microbiota besides the innate immune system, particularly acting as a fast response to potential pathogens.

## Supporting information

Intro Supplement

## Supplementary Materials

**Table S1:** Transcriptomic data.; **Table S2:** Annotation and Gene Ontology (GO) term identities of the whole transcriptome assembly.; **Table S3:** Differentially expressed genes (DE).; **Table S4:** Gene Ontology (GO) enrichment analysis of the differentially expressed genes.; **Table S5:** Assembly information of function-based identified QQ-ORFs of *A. aurita*.

## Author Contributions

Conceptualization, R. A. S. and N. W.-B.; methodology, D. L.; investigation, D. L. and N. W.-B.; formal analysis, N. W.-B.; bioinformatics analysis, V. K.; data curation, V. K.; writing—original draft preparation, N. W.-B., V. K., and R. A. S.; writing—review and editing, N. W.-B., V. K., and R. A. S., N. W.-B. and V. K.; supervision, N. W.-B. and R. A. S.; project administration, R. A. S.; funding acquisition, R. A. S.. All authors have read and agreed to the published version of the manuscript.

## Funding

This work was conducted with the financial support of the DFG-funded Collaborative Research Center CRC1182 “Origin and Function of Metaorganisms” (B2) and the DFG-funded ImmuBase project.

## Acknowledgments

We thank Sven Künzel and colleagues from the Department for Evolutionary Genetics of the Max Planck Institute for Evolutionary Biology for transcriptome sequencing.

## Conflicts of Interest

The authors declare no conflict of interest. The funders had no role in the design of the study, in the collection, analyses, or interpretation of data, in the writing of the manuscript, or in the decision to publish the results.

## Notes

### Competing Interest Statement

The authors have declared no competing interest.

