## Supplementary material for "First insights into the *Aurelia aurita* transcriptome response upon manipulation of its microbiome": Intro Supplement

**Table S1: Transcriptomic data.** (***A***) Number of raw reads and trimmed reads after filtering of native, sterile, and bacteria-challenged *Aurelia aurita* polyp pools. of under the different challenges. (***B***) Statistical data of the transcriptome assembly and the BLAST hits.

(***A***)

| treatment | raw reads | filtered reads |
|---|---|---|
| native_1 | 15,410,175 | 13,701,069 |
| native_2 | 15,248,818 | 13,492,120 |
| *Klebsiella oxytoca*_1 | 12,989,916 | 11,351,244 |
| *Klebsiella oxytoca*_2 | 11,099,887 | 9,576,528 |
| *Vibrio anguillarum*_1 | 13,413,721 | 11,957,861 |
| *Vibrio anguillarum*_2 | 15,393,642 | 13,528,712 |
| *Pseudoalteromonas espejiana*_1 | 40,705,118 | 35,719,145 |
| *Pseudoalteromonas espejiana*_2 | 16,786,302 | 14,696,662 |
| AB-treated_1 | 6,873,558 | 5,320,279 |
| AB-treated_2 | 11,639,511 | 10,234,034 |
| AB-treated_3 | 7,708,609 | 6,168,876 |
| Total | **167,269,257** | **145,746,530** |

(***B***)

| Counts of transcripts | |
|---|---|
| Total trinity 'genes' | **160,700** |
| Total trinity transcripts | **213,897** |
| GC content in % | **40** |
| **Statistics based on ALL transcripts** | |
| N10 | 3,574 |
| N20 | 2,599 |
| N30 | 2,003 |
| N40 | 1,554 |
| N50 | **1,170** |
| Median contig length | 390 |
| Average contig length | 710 |
| Total assembled bases | **151,804,471** |

| N Transcripts with BLAST ID against Swiss-Prot | |
|---|---|
| Total | |
| % | |
| Overall alignment rate in % | **94.25** |
| BUSCO Metazoa in % (n = 978) | |
| complete | **96.42** |
| single | 21.70 |
| duplicated | 74.70 |
| fragmented | 0.90 |
| missing | 2.70 |

**Table S2: Annotation and Gene Ontology (GO) term identities of the whole transcriptome assembly.**

**TableS2.xlsx**

**Table S3: Differentially expressed genes (DE).** Number of differentially expressed genes between the pairwise comparisons with their annotations and GO terms. (***Sample sheet A***) Number of genes that are up- or downregulated in AB-treated vs. native polyps. (***Sample sheet B - D***) Number of genes up- or downregulated in native polyps challenged for 30 min with $10^8$ cells/mL of (***Sample sheet B***) *Klebsiella oxytoca*, (***Sample sheet C)*** *Vibrio anguillarum*, (***Sample sheet D***) *Pseudoalteromonas espejiana.* (***Sample sheet E - H***) Number of genes exclusively upregulated in native polyps challenged for 30 min with $10^8$ cells/mL of (***Sample sheet E***) *Klebsiella oxytoca*, (***Sample sheet F***) *Vibrio anguillarum*, (***Sample sheet G***) *Pseudoalteromonas espejiana* compared to native polyps.

TableS3.xlsx

**Table S4: Gene Ontology (GO) enrichment analysis of the differentially expressed genes.** (***Sample sheet A***) enriched GO categories of upregulated genes in AB-treated polyps when compared to native polyps. (***Sample sheet B***) enriched GO categories of commonly upregulated genes in native polyps challenged for 30 min with $10^8$ cells/mL of bacteria compared to native polyps. (***Sample sheet C-E***) enriched GO categories of exclusively upregulated genes in native polyps challenged for 30 min with $10^8$ cells/mL of (***Sample sheet C***) *Klebsiella oxytoca*, (***Sample sheet D***) *Vibrio anguillarum*, (***Sample sheet E***) *Pseudoalteromonas espejiana* compared to native polyps.

TableS4.xlsx

**Table S5: Assembly information of function-based identified QQ-ORFs of *A. aurita*.**

TableS5.xlsx
